# Individual and concurrent effects of drought and chilling stresses on morpho-physiological characteristics and oxidative metabolism of maize cultivars

**DOI:** 10.1101/829309

**Authors:** Hafiz Athar Hussain, Shengnan Men, Saddam Hussain, Umair Ashraf, Qingwen Zhang, Shakeel Ahmad Anjum, Iftikhar Ali, Longchang Wang

## Abstract

Maize belongs to tropical environment and is extremely sensitive to drought and chilling stress, particularly at early developmental stages. The present study investigated the individual and combine effects of drought (15% PEG-Solution) and chilling stress (15°C/12°C) on the morpho-physiological growth, osmolyte accumulation, production and regulations of reactive oxygen species (ROS) and the activities of antioxidants in two maize hybrids i.e., ‘XD889’ and ‘XD319’ and two inbred cultivars i.e., ‘Yu13’ and ‘Yu37’. Individual and combined drought and chilling stresses stimulated the production of O2^□^, H_2_O_2_, OH^□^ and enhanced malondialdehyde (MDA) contents which led to reduced photosynthetic pigments and morphological growth. Drought, chilling and drought + chilling stress conditions induced the compatible osmolytes, ROS detoxifying proteins and antioxidants to counterbalanced the oxidative damage. It was found that the concurrent occurrence of drought + chilling stress was more lethal for maize seedling growth than the drought and chilling individual stresses. However, the performance of hybrid maize cultivars (XD889 and XD319) was better than the inbred maize cultivars (Yu13 and Yu37). For improving tolerance to individual and concurrent drought and chilling stress in maize, future research focus should be on developing genetically engineer plants that have the ability to generate specific response against sub-optimal temperature and water deficit conditions.

## Introduction

Crop plants are often exposed to multiple abiotic stresses simultaneously of which concurrent effects of sub-optimal temperature and moisture deficit conditions are perhaps the most deleterious for plant growth [1,2,3]. Drought and chilling stresses cause substantial reduction in growth and yield attributes of plants by disturbing normal plant metabolism [2,4]. For instance, the sub-optimal temperature causes water deficiency in maize seedlings by substantial reduction in root water uptake coupled with leaves transpiration [5]. Conversely, drought-pretreatment induces the stomatal closure and decline the transpiration in maize [6]. Furthermore, Hussain et al., (2018) indicated that the drought and chilling stresses triggered the plant-water relations but these effects are possibly more detrimental under the simultaneous occurrence of these both factors [2]. However, the destructive effects of low temperature and water deficiency on the crop plants depend on the duration, nature and severity of stress and the plant developmental stages. For exploring the nutrients and water, suboptimal temperature decreased the root growth of maize seedlings by decreasing the length and biomass of the roots [7], in contrast, drought stress increased the root length of sunflower plants but later on decreased with the increasing of drought period [8]. Previous findings have showed that the osmotic balance of the plants is more affected by drought as compared to chilling stress [9].

Generally, the effects of both these stresses on plant morphological traits are quite similar, however their mechanisms to affect the physio-biochemical processes of plants may vary [10]. Most often, individual and/or concurrent drought and chilling stresses results in excessive production of reactive oxygen species (ROS) that cause structural and functional damage to cell and its constituent [2,11], whereas the degree of oxidative damage may be accelerated synergistically under combined drought and low temperature conditions [12,13,14].

Osmo-regulation by accumulation of different osmolytes plays significant role in turgor maintenance and protection of macromolecules in dehydrating cells under stressful conditions [15,16]. In order to control the overproduction of ROS, the plants have a highly efficient and sophisticated enzymatic and non-enzymatic antioxidative defense system [1,18,19]. Guo et al. (2006) reported that drought and chilling tolerance in plants is strongly linked with the enhanced activities/levels of antioxidants under stress condition [20].

Maize is a thermophilic crop and belongs to the tropical environment with optimum growth temperature around 28°C [21]. Nevertheless, at emergence and early growth stages, the maize seedling are extremely sensitive to drought and low temperature. Water deficiency and sub-optimal temperatures (<12-15°C) during seedling growth can be detrimental to subsequent crop productivity [4,6]. Although the inhibitory effects of sub-optimal temperature and low moisture supply on morpho-physiological growth and productivity of various crops previously reported [20,22,23,24], however, little information is available about the individual and concurrent effect of sub-optimal temperature and low moisture supply on morpho-physiological growth and oxidative metabolism in hybrid and inbred maize cultivars. Therefore, present study was conducted to assess the morpho-physiological changes, extent of ROS production and regulations in the activities of antioxidants in maize seedlings under the influence of individual and combined sub-optimal temperature and low moisture supply.

## Materials and Methods

### Plant Material and Growth Conditions

This experiment was conducted in the controlled growth chamber by using hydroponic solution at the College of Agronomy and Biotechnology (CAB), Southwest University (SWU), Chongqing, China (longitude 106º 26’ 02’’E, latitude 29º 49’ 32’’ N, and altitude 220 m) during spring 2016. The seeds of two maize hybrids i.e., Xida889 (XD889), Xida319 (XD319) and two maize inbred i.e., Yu13 and Yu37 were obtained from Maize Research Institute, CAB, SWU, Chongqing, China. The initial germination rate and moisture (on dry weight basis) content of the seeds were >90% and <10%, respectively. Prior to sowing, seeds were surface sterilized with NaOCl solution to minimize contamination and rinsed thrice with sterile distilled water.

The maize seed were kept on wet muslin cloth in normal condition for germination. After germination (7 days after sowing; DAS), seedlings were transplanted to plastic pots (30 cm × 20 cm × 12 cm) filled with modified Hoagland’s nutrient solutions [25]. Plastic pots containing 4 L of respective solution and a floating board with four separated sections (for maize cultivars), were used. Six seeds of each cultivar were sown. Six plants per variety were transplanted in each section of the board. The experiment was laid out in a completely randomized design with three replicates.

### Stress Treatments

After transplanting, the seedlings were subjected to drought and/or low temperature stresses. For control and drought stress treatments, temperature was set as, 25°C at day and 20°C at night with 12h interval. For chilling and chilling + drought stresses treatments, temperature was set as 15°C at day and 12°C at night with 12h interval. Drought stress was imposed by using 15% polyethylene glycol-6000 (PEG) solution. The combined stress consisted of simultaneous treatment with PEG and chilling stress. Nutrient solution with or without PEG (15%) was changed after every 4 days. After 23 DAS, seedlings were harvested and growth parameters were recorded whereas fresh maize seedlings were stored at −80°C for measuring different biochemical analysis.

### Observations

#### Measurement of Growth Parameters

Shoot and root length of maize seedlings was recorded with a meter scale while an electronic weighing balance was used to measure the shoot and root fresh biomass. The maximum leaf width and stem diameter were measured with Digimatic caliper (500-197-30, Mitutoyo group, Japan), while number of leaves was counted manually. Total root length, root surface area, average diameter and root volume were analyzed with WinRHIZO by using Epson Perfection V700 Photo Flatbed scanner (B11B178023, Indonesia).

#### Photosynthetic pigments

Chlorophyll (Chl a, Chl b and total Chl) concentration were determined according to Peng and Liu (1992) method [26]. Extraction of 250 mg leaf without vein (leaf blade) was done with 10 ml ethanol-acetone (vol. 1:2), and the extract was transferred to 15 ml tube. The tubes were placed in dark to avoid light for 24 hours. The absorbance was measured at 645 nm, 663 nm, and 652 nm. The chlorophyll content was computed by the following formulae:

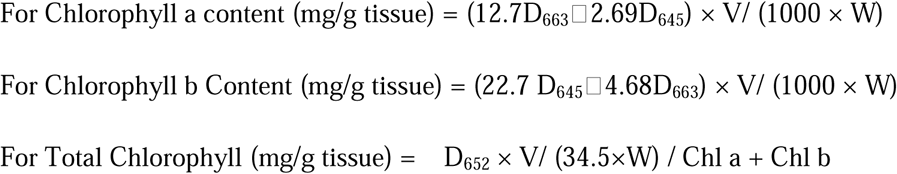

Where, D663, D645 and D652 respectively are the corresponding wavelengths of the light density value, V is extracting liquid volume and W is leaf fresh weight.

#### Lipid Peroxidation and ROS Activity

Lipid peroxidation was determined as malondialdehyde (MDA) content, measured by thiobarbituric (TBA) method using ‘MDA Detection Kit (A003)’ as described previously [27]. The absorbance for MDA was measured at 532 nm and expressed as nmol/g fresh weight.

The contents of hydrogen peroxide (H_2_O_2_), hydroxyl free radical (OH^−^) and superoxide anion radical (O^−2^) in the leaves of maize seedlings were determined using the commercial ‘H_2_O_2_ Detection Kit (A064)’, ‘OH^−^ Detection kit (A018)’ and ‘O^−2^ Detection kit (A052)’, respectively, obtained from Nanjing Jiancheng Bioengineering Institute, China. The H_2_O_2_ bound with molybdenic acid to form a complex, which was measured at 405 nm and the content of H_2_O_2_ was then calculated. The OH^−^ was expressed as unit mg^-1^ protein, and one unit was the amount required to reduce 1 mmol/L of H_2_O_2_ in the reaction mixture per minute at 37°C. The O^−2^ inhibition per g tissue protein for 40 minutes in 37°C reaction which equals to superoxide anion radical inhibition caused by 1mg Vc is considered as 1 anti-superoxide anion radical activity unit.

#### Estimation of enzymatic and non-enzymatic antioxidants

The activities of enzymatic antioxidants were detected by using the commercial kits in accordance with the manufacturer’s instructions as described earlier [27, 28]. The kits for superoxide dismutase (A001), peroxidase (A084-3), catalase (A007-2), glutathione peroxidase (A005), glutathione reductase (A062), and glutathione S-transferase (A004) were purchased from the same company as mentioned above. The absorbance readings of SOD, POD, CAT, GPX, GR, and GST were detected at 550 nm, 420 nm, 405 nm, 412 nm, 340 nm, and 412 nm, respectively (Tecan-infinite M200, Switzerland). The SOD, POD, CAT, GPX, and GST activities were expressed as units U/mg protein, while GR activity was demonstrated as units U/g protein. The units of the antioxidant enzyme activities were defined as follows: one unit of SOD activity was the amount of enzyme required to decrease the reference rate to 50% of maximum inhibition; one unit of POD activity was defined as the amount of enzyme necessary for the decomposition of 1 µg substrate in 1 min at 37°C; one unit of CAT activity was defined as the amount of enzyme required to decompose the 1 µM H_2_O_2_ in 1 second at 37^°^C; One unit is GPX activity was the amount of enzyme required to oxidize 1 µM GSH in 1 minute at 37°C; one unit of GR activity was defined as the amount of enzyme depleting 1 mM NADPH in 1 min; and one unit of GST activity was defined as the amount of enzyme depleting 1 µM GSH in 1 min.

The glutathione (GSH), ascorbic acid (Vitamin C), and vitamin E content in leaves of maize seedlings were measured by colorimetric method using the ‘GSH (deproteinization) assay kit-A006’, ‘Vitamin C assay kit-A009’, and ‘Vitamin E assay kit-A008’, respectively. The absorbance for GSH, Vc, and Ve were recorded at 420 nm, 536 nm, and 533 nm (Tecan-infinite M200, Swit), respectively. The GSH content were expressed as mg/g protein, the Vc and Ve expressed as µg g^− 1^ tissue fresh weight. The commercial ‘kit-A015’ was used for determination of total antioxidant capability (T-AOC). The absorbance for T-AOC was measured at 520 nm and data were expressed as U/mg protein. One unit of T-AOC was defined as the amount that increased the OD value per mg tissue protein per minute by 0.01 reaction system at 37°C.

#### Assay of Osmolyte Accumulation Profiles

Free proline (FP) contents were assessed by following the method of Shan et al (2007) [29]. Briefly, fresh leaves (0.5 g) was extracted with 5 ml of 3% sulphosalicylic acid at 100 °C for 10 min with shaking. The extracts were filtered through glass wool and analysed for proline content using the acid ninhydrin method and the absorbance was read at 520 nm. The resulting values were compared with a standard curve constructed using known amounts of proline (Sigma, St Louis, MO, USA).

Total soluble sugar was estimated by anthracene ketone method as described by Zong and Wang (2011) [30]. Fresh leaves (0.2 g) was homogenized with 25 ml distilled water and centrifuged at 4000 rpm for 20 minutes. Anthracene (0.1 g) was dissolved in 100 ml diluted sulfuric acid. One ml extract and 5 ml anthracene sulfuric acid reagent were taken in a tube, shaken and put in boiling bath for 10 minutes. The mixture was kept at room temperature for stabilizing and the absorbance was read at 620nm. The content of total protein in the leaves of maize seedling was determined by Coomassie brilliant blue method using ‘Total Protein Quantification Kit (A045-2)’ obtained from Nanjing Jiancheng Bioengineering Institute, China and unit of protein was quantified as g/L sample solution. Total amino acid in maize seedlings was determined by using ‘(T-AA) Detection Kit A026’ obtained from Nanjing Jiancheng Bioengineering Institute, China. The content of total amino acid was expressed as µmol/g fresh weight.

#### Statistical Analysis

The data collected were statistically analyzed following the analysis of variance technique using Statistix 8.1 (Analytical Software, Tallahassee, FL, USA) software and the mean variance of the data was analyzed using the least significant difference (LSD) test at the 0.05 probability level. Sigma Plot 12.5 (Systat Software Inc., San Jose, CA, USA) was used for graphical presentation of the data.

## Results

### Shoot Growth

Pronounce influence were observed in the maize shoot growth under the drought, chilling and combination of drought +chilling stresses (Figure 1 and 2). Our results indicated that shoot length, shoot fresh weight and stem diameter of all four maize cultivars was considerably reduced under stress conditions as compared with control, nevertheless, the negative effects of combined drought +chilling stresses were more severe for all these traits than their individual effects (Figure 1). However, the shoot lengths of XD889 and Yu37 cultivars under drought stress were statistically similar (*P*□0.05) with control (no drought+ normal temperature) treatment (Figure 1a). Compared with control, the significant reductions in shoot fresh weight were noticed under drought, chilling and drought + chilling stress conditions, respectively. Furthermore, the influence of chilling stress on stem diameter of Yu13 and Yu 37 was non-significant (P□0.05) compared with control (Figure 1c). The shoot growth performance of hybrid maize cultivars was better than inbred cultivars under control as well as stress conditions.

**Figure 1:**
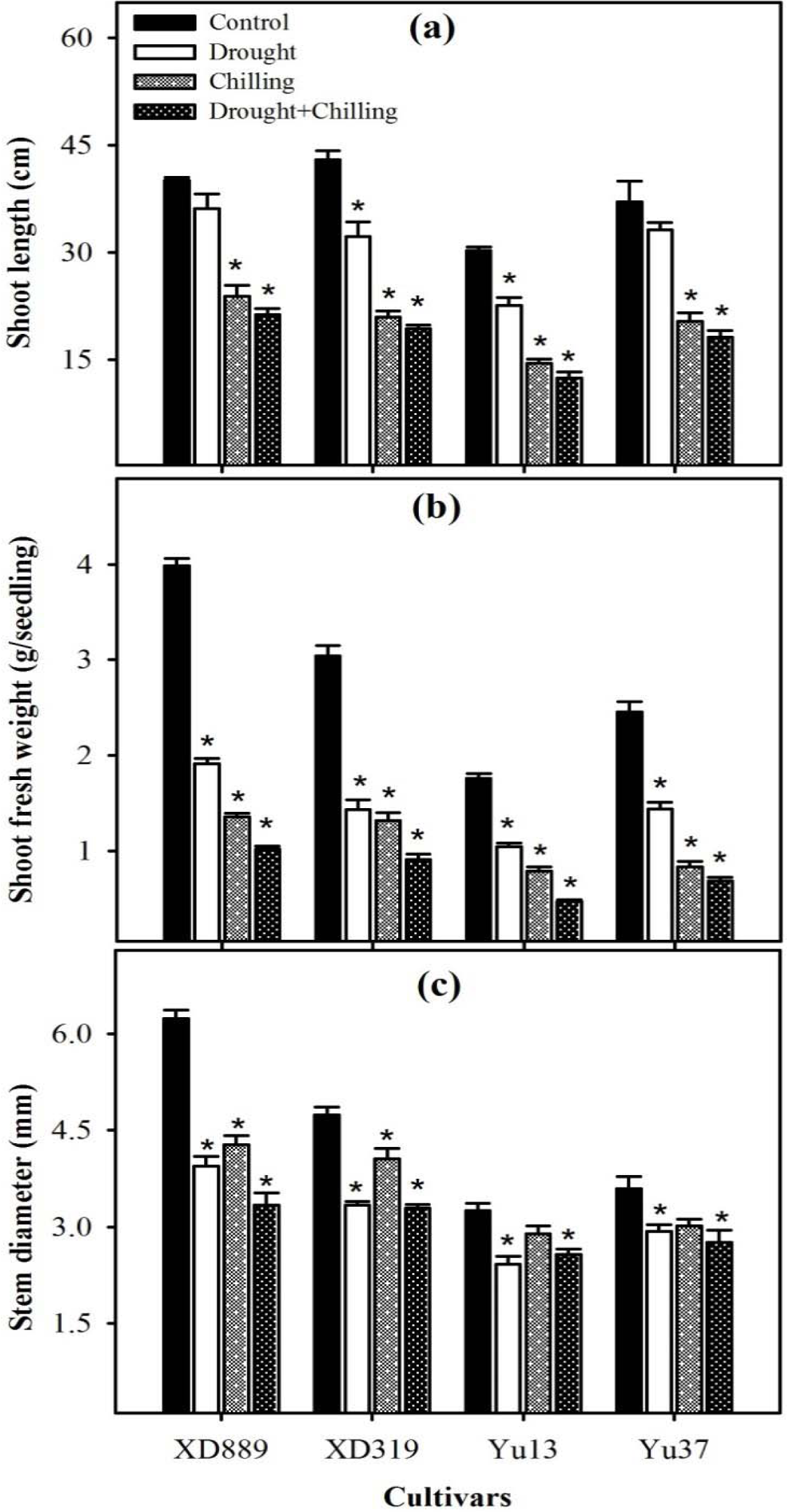
Shoot length (a), shoot fresh weight (b), and stem diameter (c) of four maize cultivars as influenced by drought, chilling and drought + chilling stress. Vertical bars above mean indicate standard error of three replicates. Mean value for each treatment with * indicate significant differences by the LSD-test (P ≤ 0.05).

**Figure 2:**
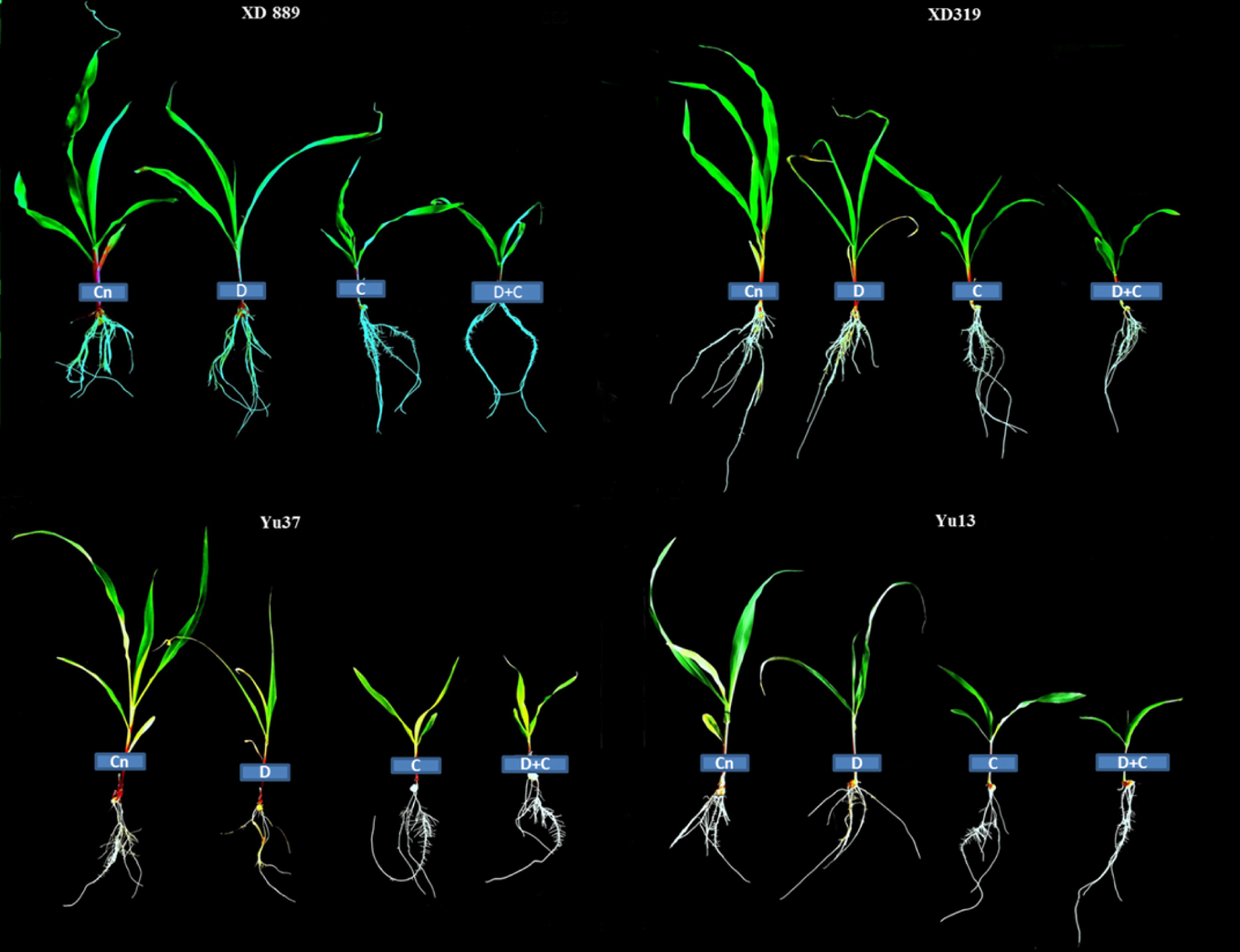
Individual and interactive effect of drought and chilling stresses on the seedlings growth of maize (cv. XD889, XD319, Yu13, Yu37). Cn: control, D: drought, C: Chilling, D+C: drought + chilling.

### Leaf Growth

Number of leaves and leaf width of four maize cultivars was noticeably decreased under stress conditions as compared with control, on the other hand, the negative effects of drought + chilling stresses was more severe for all maize cultivars then their individual effects (Figure 2 and 3). However, the number of leaves of XD889 cultivar under chilling and drought + chilling was significantly affected as compared with control. Influence of drought stress on leaf width of XD319 and Yu37 cultivars was statistically similar (*P*□0.05) with control. Overall, the leaf growth of hybrid maize cultivars was better than inbred cultivars under control as well as stress conditions (Figure 3).

**Figure 3:**
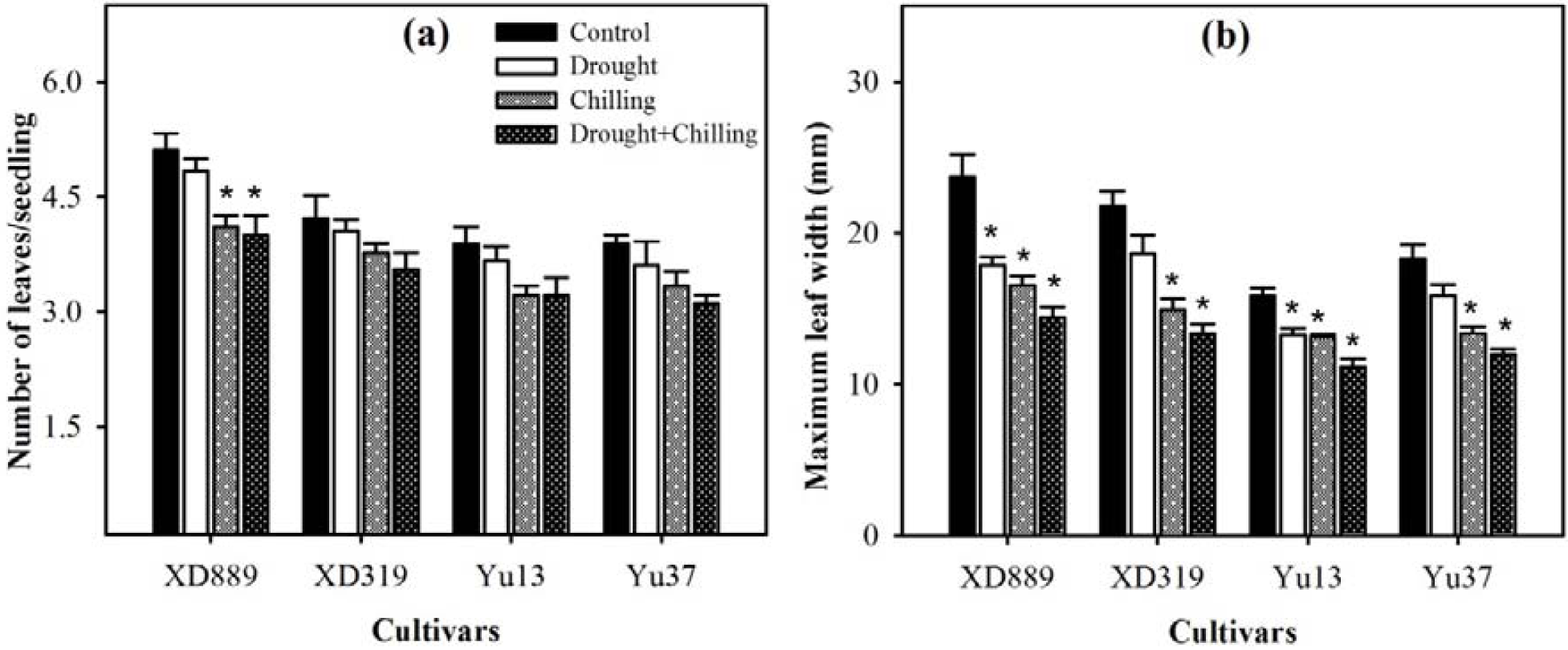
No of leaves (a) and maximum leaf width (b) of four maize cultivars as influenced by the drought, chilling and drought + chilling stress. Vertical bars above mean indicate standard error of three replicates. Mean value for each treatment with * indicate significant differences by the LSD-test (P ≤ 0.05).

### Root Growth

Individual and concurrent drought and chilling stress conditions severely affected the maize root growth (Figure 4, 5 and 6). Root length of XD889 increased under drought stress but reduced under chilling and drought + chilling stress conditions. Likewise, root length of XD319, Yu13 and Yu37 reduced under respective stress conditions as compared with control. However, the root length of XD319 under drought + chilling stress, Yu13 under drought, chilling and drought + chilling stress, Yu37 under chilling and drought + chilling stress were significantly affected as compared with control (Figure 4a). Root fresh weight of all maize cultivars were decreased under respective stress conditions, however, the influence of drought + chilling stress on root fresh weight of Yu-37 was statistically similar (*P*□0.05) with control (Figure 4b).

**Figure 4:**
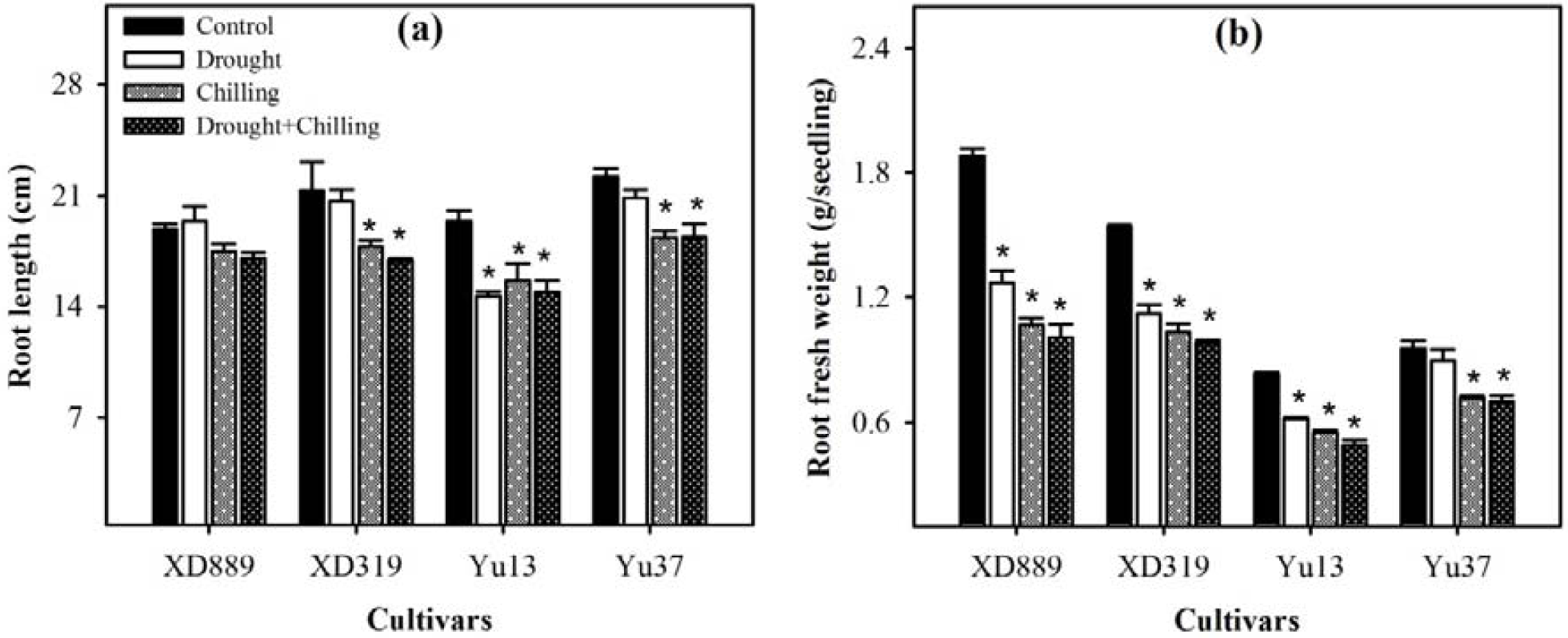
Root length (a) and root fresh weight (b) of four maize cultivars as influenced by the drought, chilling and drought + chilling stress. Vertical bars above mean indicate standard error of three replicates. Mean value for each treatment with * indicate significant differences by the LSD-test (P ≤ 0.05).

**Figure 5:**
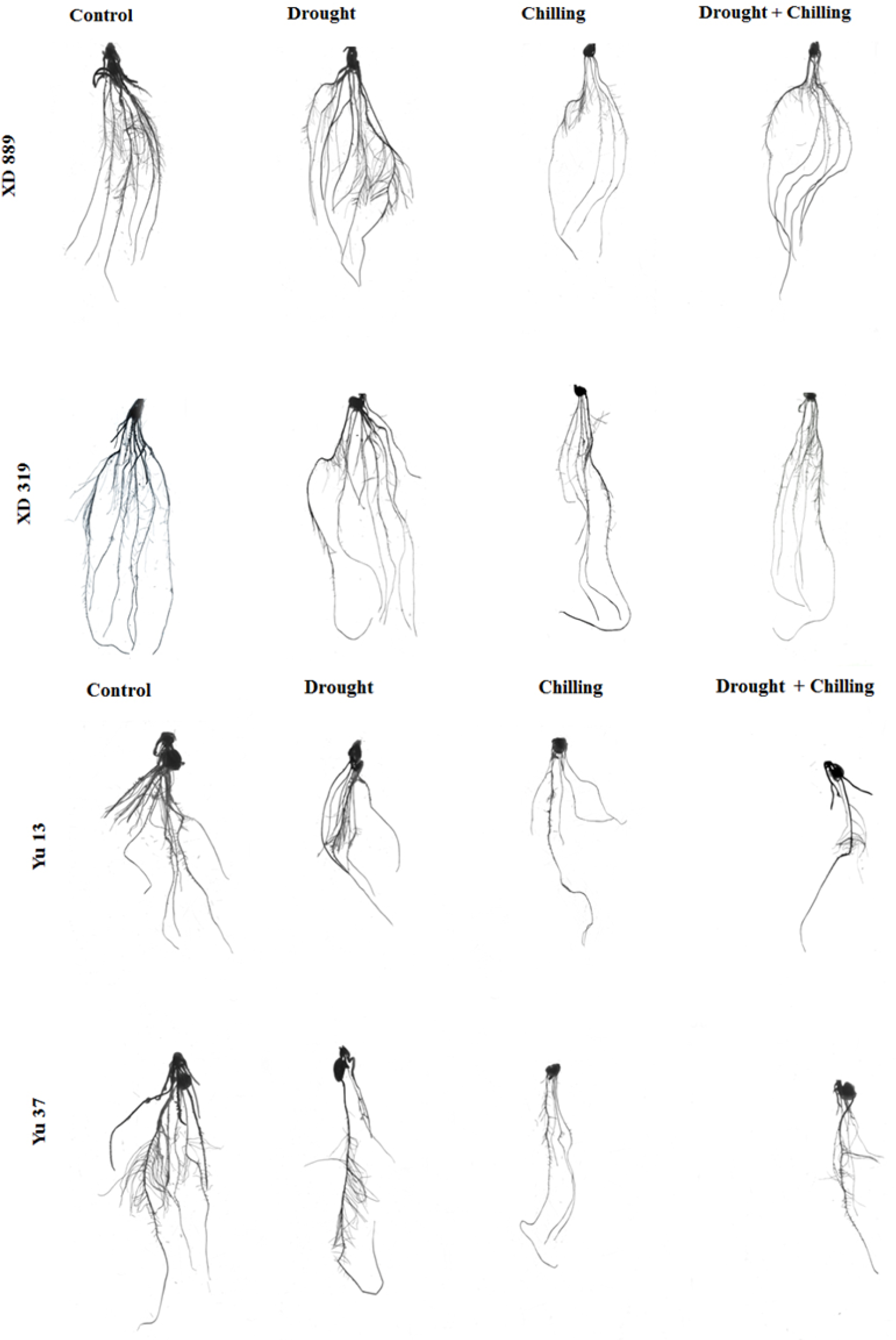
Root growth of maize seedlings (cv. XD889, XD319, Yu13, Yu37) under drought, chilling and drought + chilling stress conditions.

**Figure 6:**
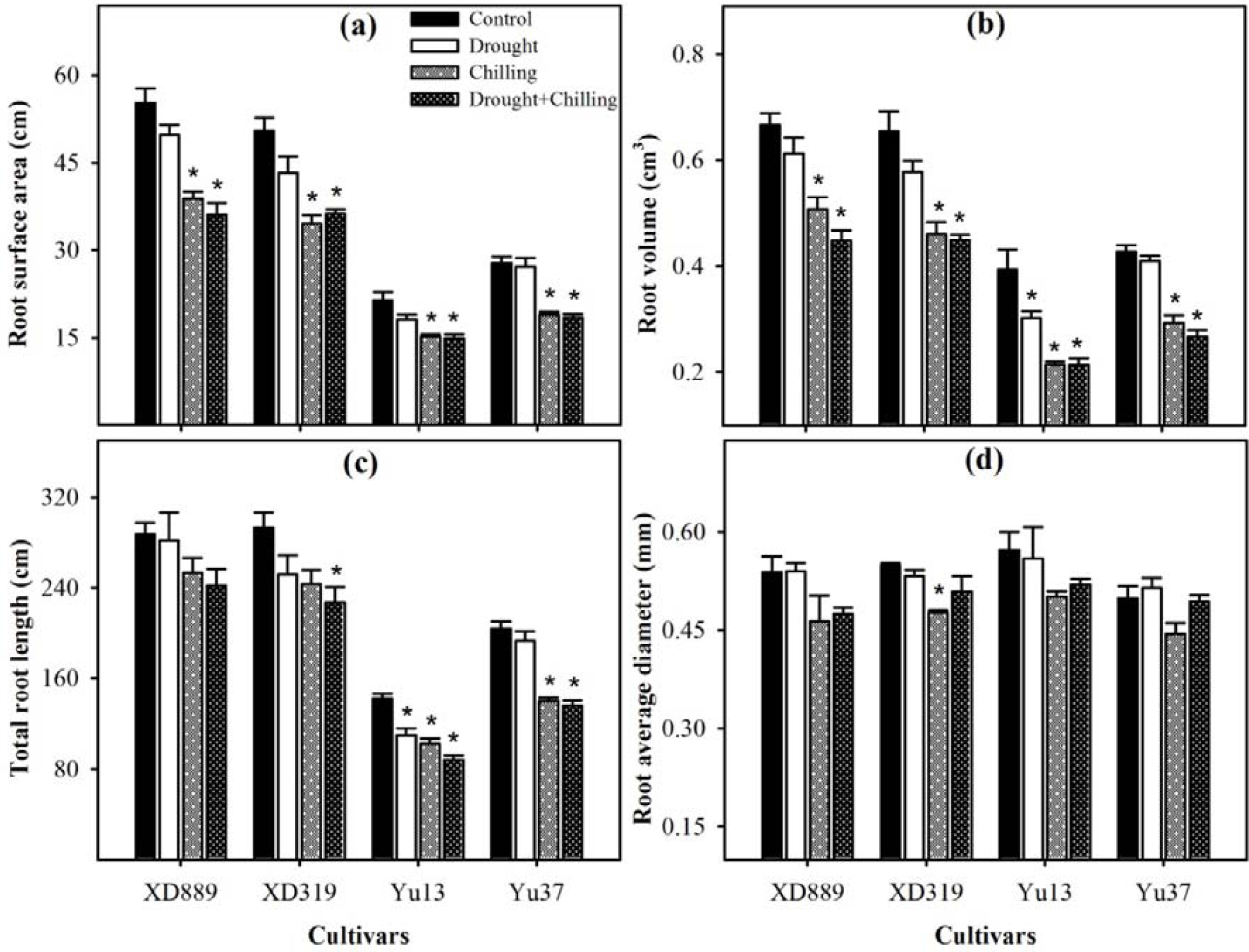
Root surface area (a), root volume (b), total root length (c), and root average diameter (d) of four maize cultivars as influenced by the drought, chilling and drought + chilling stress. Vertical bars above mean indicate standard error of three replicates. Mean value for each treatment with * indicate significant differences by the LSD-test (P ≤ 0.05).

As compared with control, root surface area, root volume and total root length of all maize cultivars were decreased under respective stress conditions. Furthermore, impact of drought stress on the root surface area of all maize cultivars and root volume of XD889, XD319 and Yu37 was statistically similar (P□0.05) as control treatment. However, total root length of XD319 under drought + chilling stress, Yu13 under all stress conditions and Yu37 under chilling and drought + chilling stress conditions, were significantly declined as compared with control treatment. Root average diameter of XD319 and reduced Yu13 under respective stress conditions as compared with control. Conversely, the root average diameter of XD889 and Yu37 increased by drought stress but reduced under chilling stress and drought +chilling stress respectively (Figure 6d). Overall the root growth performance of maize hybrid cultivars was betters then the maize inbred cultivars under control as well as stress conditions (Figure 5 and 6).

### Photosynthetic pigments

The leaf chlorophyll (Chl) concentration of maize cultivars hampered under stress conditions as compared with control, whereas, the negative effects of drought stress were more severe for chlorophyll contents than the individual effects of chilling and drought + chilling stresses (Figure 7). However, the impact of drought and drought + chilling stress on the Chl a of XD 889 and Yu13 was significant as compared with control (Figure 7a). Chl b decreased in XD889, XD319 and Yu13 under respective stress conditions. While, the Chl b contents were increased in Yu37 by chilling stress and then decreased by drought and drought + chilling stress conditions compared with control. Furthermore, drought, chilling and drought + chilling stress conditions reduced the total chlorophyll in XD319, Yu13 and Yu37 while, increased in XD 889 under chilling, and then decreased under drought and drought + chilling stress conditions as compared with control (Figure 7c).

**Figure 7:**
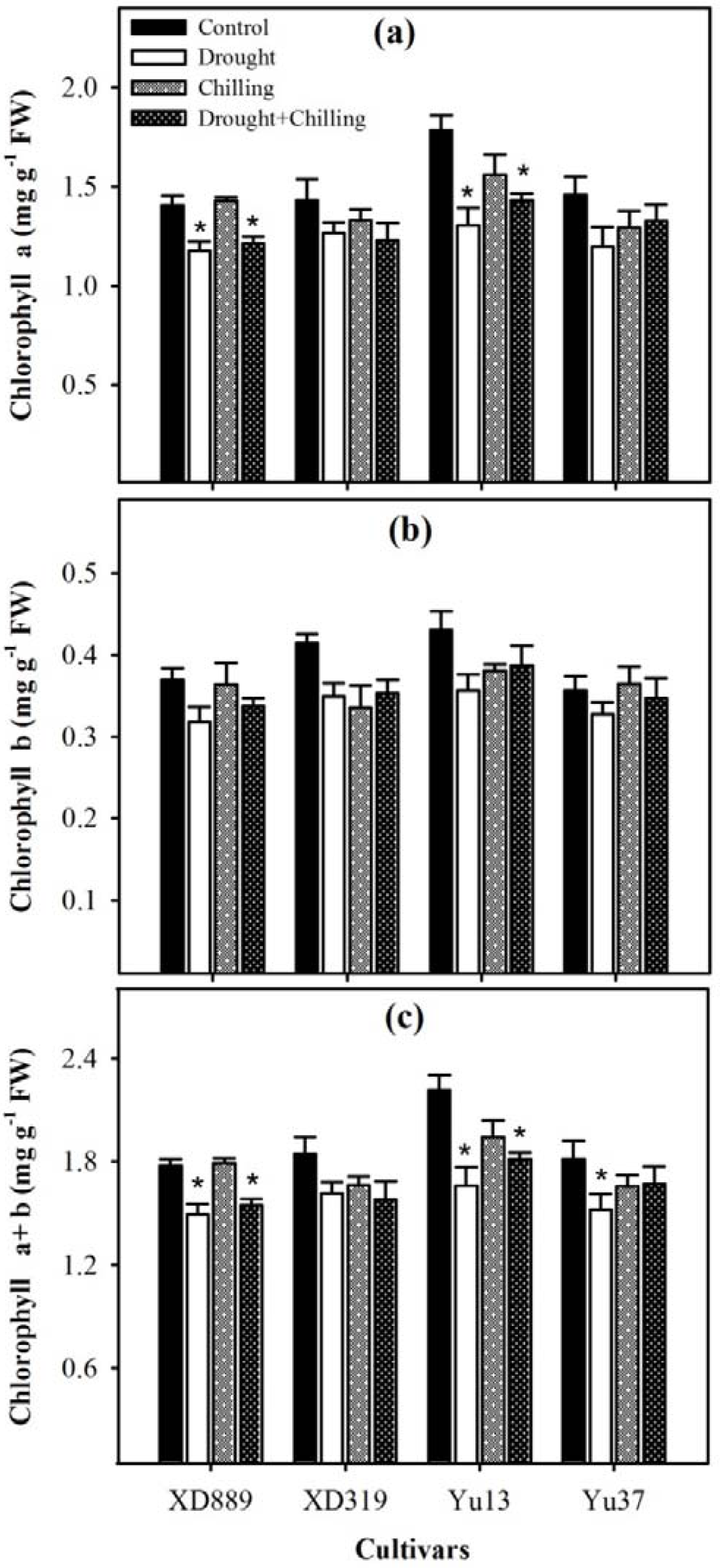
Chlorophyll a content (a), chlorophyll b content (b), chlorophyll a+b content (c) in four maize cultivars as influenced by the drought, chilling and drought + chilling stress. Vertical bars above mean indicate standard error of three replicates. Mean value for each treatment with * indicate significant differences by the LSD-test (P ≤ 0.05).

### Reactive oxygen species and lipid peroxidation

Production of reactive oxygen species (ROS) at normal level is essential and beneficial for the physiological functions of the plants but over accumulation of ROS caused oxidative stress which is destructive for biological process in plants. The level of Superoxide anion (O_2_^-^), Hydrogen peroxide (H_2_O_2_), hydroxyl radical (OH^-^), and lipid peroxidation in term of MDA production in all maize cultivars were significantly increased under individual and combine drought and chilling stresses (Figure 8). The higher concentrations of ROS as a result of these stress conditions caused oxidative damage and reduced lipid peroxidation in term of MDA, and lead to the oxidative destruction of the plants cell which stunted the overall plants growth.

**Figure 8:**
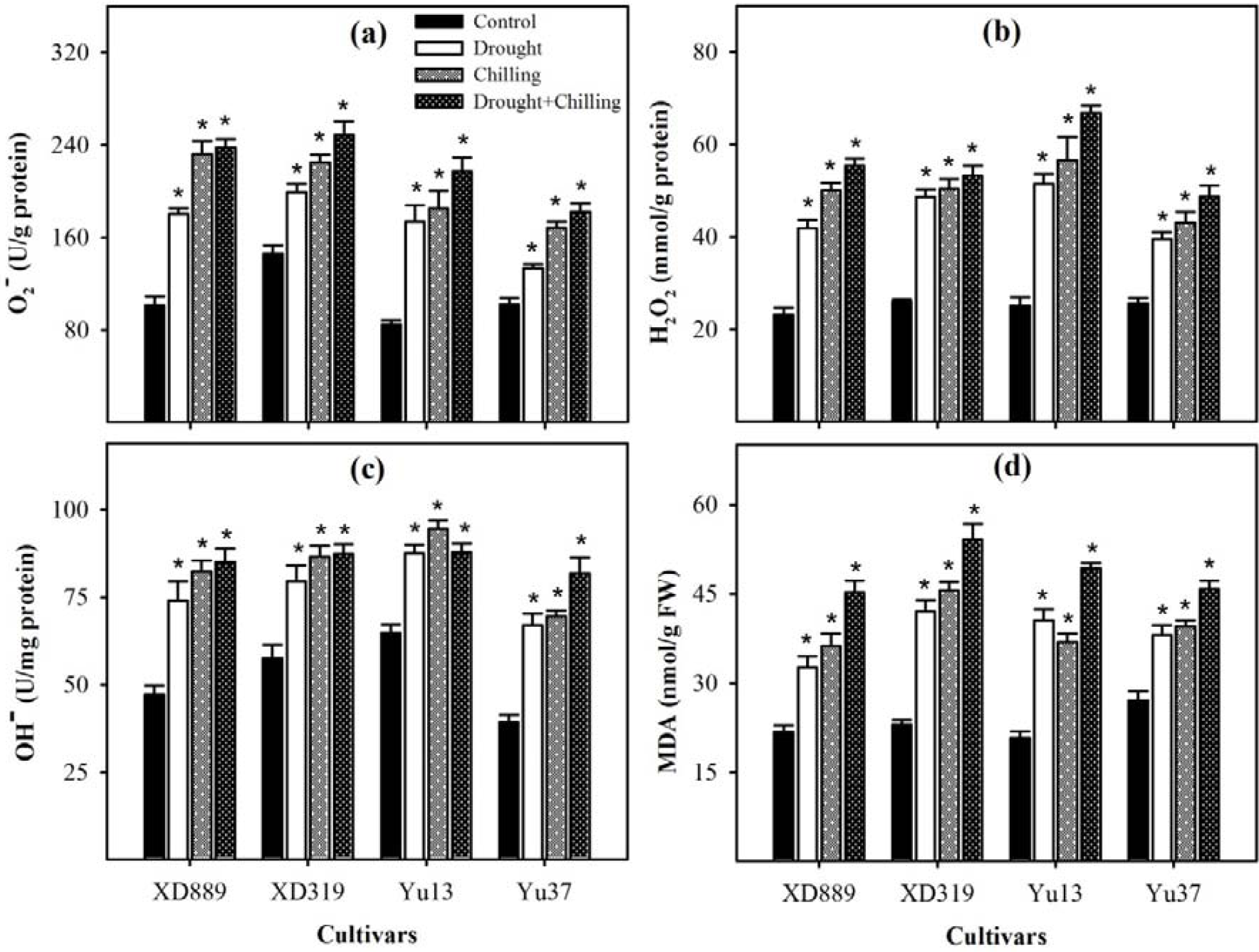
Accumulation of superoxide radical (a), hydrogen peroxide (b), hydroxyl ion (c), and malondialdehyde content (d) in the leaves of four maize cultivars as influenced by the drought, chilling and drought + chilling stress. Vertical bars above mean indicate standard error of three replicates. Mean value for each treatment with * indicate significant differences by the LSD-test (P ≤ 0.05).

### Enzymatic antioxidants

SOD reduced in all maize cultivars under drought, chilling and drought + chilling stress treatments, however, the impact of drought stress on SOD of XD889 and Yu37 was statistically similar (P□0.05) with control (Figure 9). Compared with control, CAT substantially increased in XD889, drought stress reduced CAT in XD319 but increased under chilling stress and significantly improved under combine stress. The CAT activity was reduced in Yu13 but increased in Yu37 under respective stress conditions (Figure 9b). Imposition of drought, chilling and drought + chilling stress considerably reduced the POD in all cultivars as compared with control. GR activity were increased in XD889, XD319 and Yu37 decreased in Yu13 under drought, chilling and drought + chilling stress treatments, compared with control (Figure 9d).

**Figure 9:**
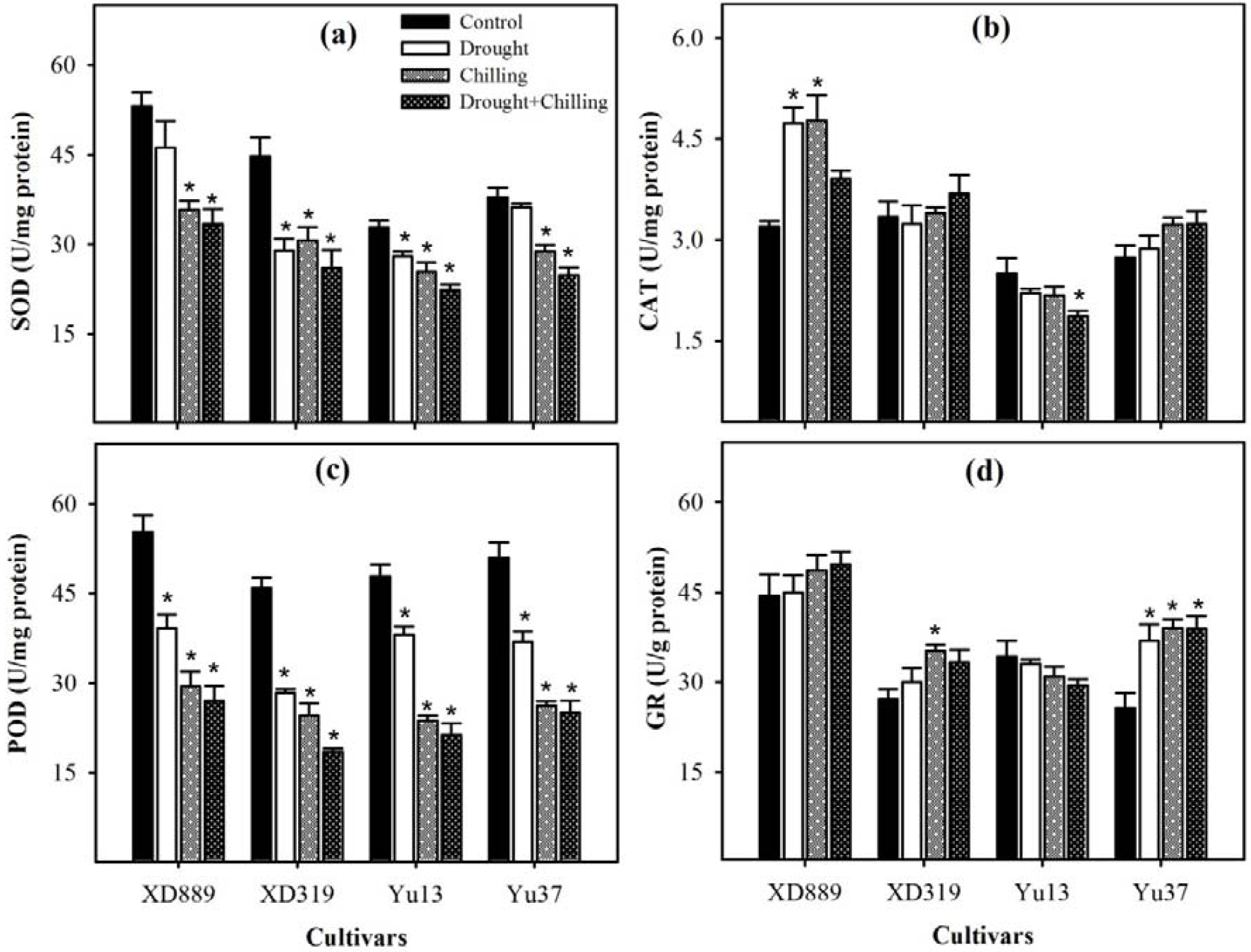
Activities of superoxide dismutase (a), catalase (b), peroxidase (c), and glutathione reductase (d) in the leaves of four maize cultivars as influenced by the drought, chilling and drought + chilling stress. Vertical bars above mean indicate standard error of three replicates. Mean value for each treatment with * indicate significant differences by the LSD-test (P ≤ 0.05).

In all maize cultivars, GSH-PX and GST noticeably increased under stress conditions as compared with control, nonetheless, the enhancement was more by drought + chilling combine stress as compared with their individual effects (Figure 10). Compared with control, the GSH-PX activity was improved in all maize cultivars under respective stresses. However, effect of drought stress on GSH-PX inYu37, and effect of drought, chilling and drought + chilling stress treatments on GSH-PX in XD 319 was statistically similar (P□0.05) as compared with control (Figure 10a). In addition, the GST activity of XD889 under drought stress, XD319 under drought and chilling stress treatments, Yu13 under drought, chilling and drought + chilling stress treatments were remained non-significant (P□0.05) than control. Overall performance of hybrid maize cultivars was better than inbred maize cultivars under stress conditions as well as control condition (Figure 10).

**Figure 10:**
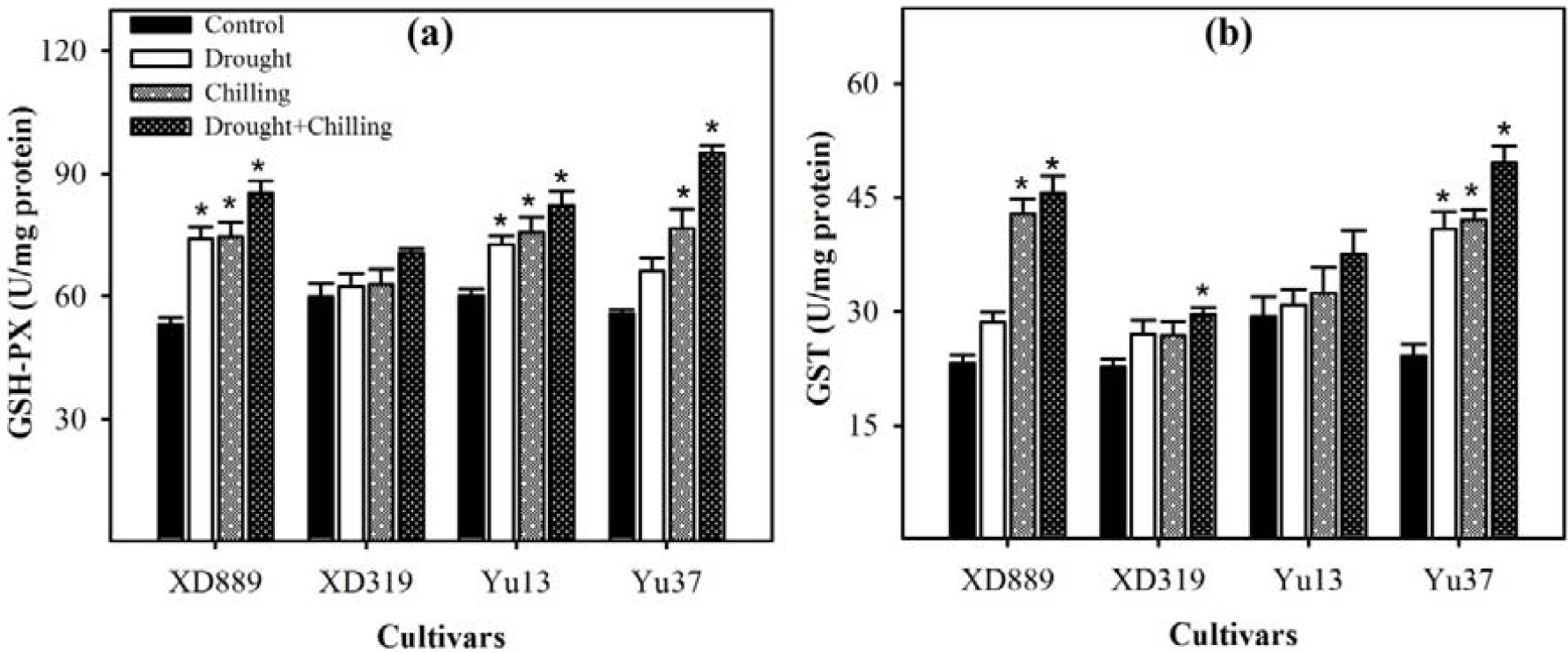
Activities of glutathione peroxidase (a), and glutathione S-transferase (b) in the leaves of four maize cultivars as influenced by the drought, chilling and drought + chilling stress. Vertical bars above mean indicate standard error of three replicates. Mean value for each treatment with * indicate significant differences by the LSD-test (P ≤ 0.05).

### Non-enzymatic antioxidants

The non-enzymatic antioxidants in the leaves of maize were triggered under drought, chilling and drought + chilling stress conditions. Results showed that glutathione content, vitamin C, vitamin E and total antioxidant enzymes severely hampered under stress treatments as compared with control (Figure 11). For instance, the GSH contents were reduced under drought stress but increased under chilling stress and drought + chilling stress in XD889 and XD319, respectively, compared with control. The GSH contents were reduced in Yu13 but increased in Yu37 under respective stress treatments. Furthermore, the effects of drought + chilling stress on GSH in XD889 were significant as compared with control (Figure 11a). Drought, chilling and drought + chilling stress conditions increased V_C_ in all maize cultivars, as compared with control. However, Vc of XD889 under chilling and drought + chilling stresses, XD319 and Yu37 under drought + chilling stress was significant as compared with control (Figure 11b). V_E_ increased in XD889, XD319 and Yu37 under respective stresses, whereas, V_E_ enhanced in Yu13 under drought stress, but reduced under chilling and drought + chilling stresses. Moreover, compared with control, the T-AOC enhanced in all maize cultivars, under drought, chilling and drought + chilling stress treatments. Additionally, influence of drought stress on T-AOC in XD319 and Yu13 was statistically similar (P□0.05) with control (Figure 11d).

**Figure 11:**
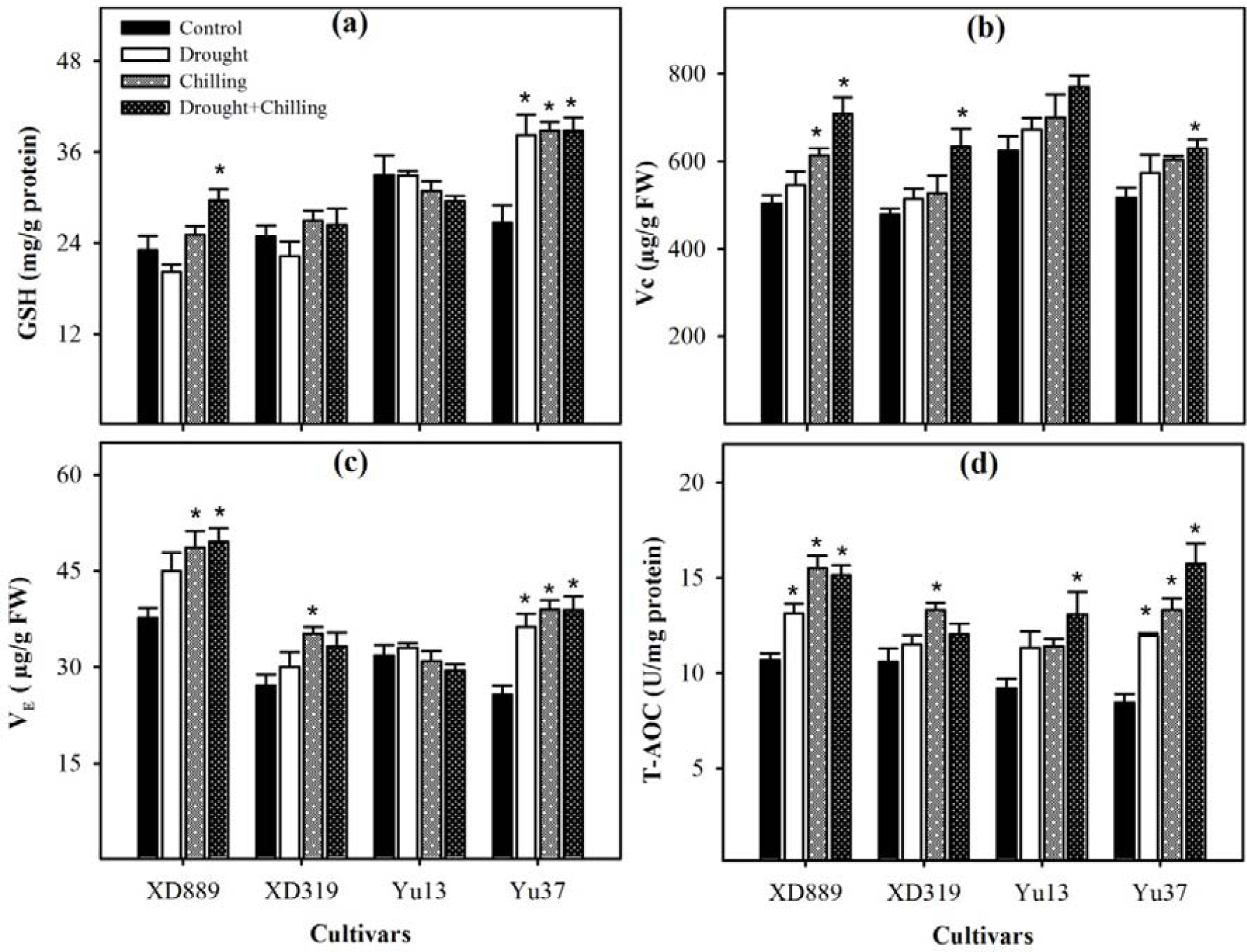
Levels of glutathione content (a), vitamin C (b), vitamin E (c), and total antioxidant capacity (d) in the leaves of four maize cultivars as influenced by the drought, chilling and drought + chilling stress. Vertical bars above mean indicate standard error of three replicates. Mean value for each treatment with * indicate significant differences by the LSD-test (P ≤ 0.05).

### Metabolites accumulation

Drought and chilling stresses regulated the accumulation protein content, free proline content, total soluble sugars and total amino acid in all maize cultivars (Figure 12). Exposure of drought, chilling and drought + chilling enhanced the protein in all maize cultivars as compared with control, however, the protein content of XD889 and Yu37 under drought stress was statistically similar (P□0.05) with control. Free proline content was found to be significantly higher in all maize cultivars under respective stresses as compared with control. Total soluble sugar reduced in XD319, Yu13, and Yu37 under respective stress treatments as compared with control, while in XD889, total soluble sugars were increased under drought stress but decreased under chilling and drought + chilling stress treatments. In addition, total soluble sugars in XD889, XD319 and Yu37 under chilling stress, Yu13 under chilling and drought + chilling stress treatments were significant as compared with control (Figure 12c). The TAA increases in XD889, XD319, Yu37 respective stress condition but in Yu13 reduced under drought stress and then increased under chilling and drought + chilling stresses as compared with control, furthermore, the influence of chilling and drought + chilling treatments on the TAA of XD889 and drought + chilling on Yu37 were significant as compared with control (Figure 12d).

**Figure 12:**
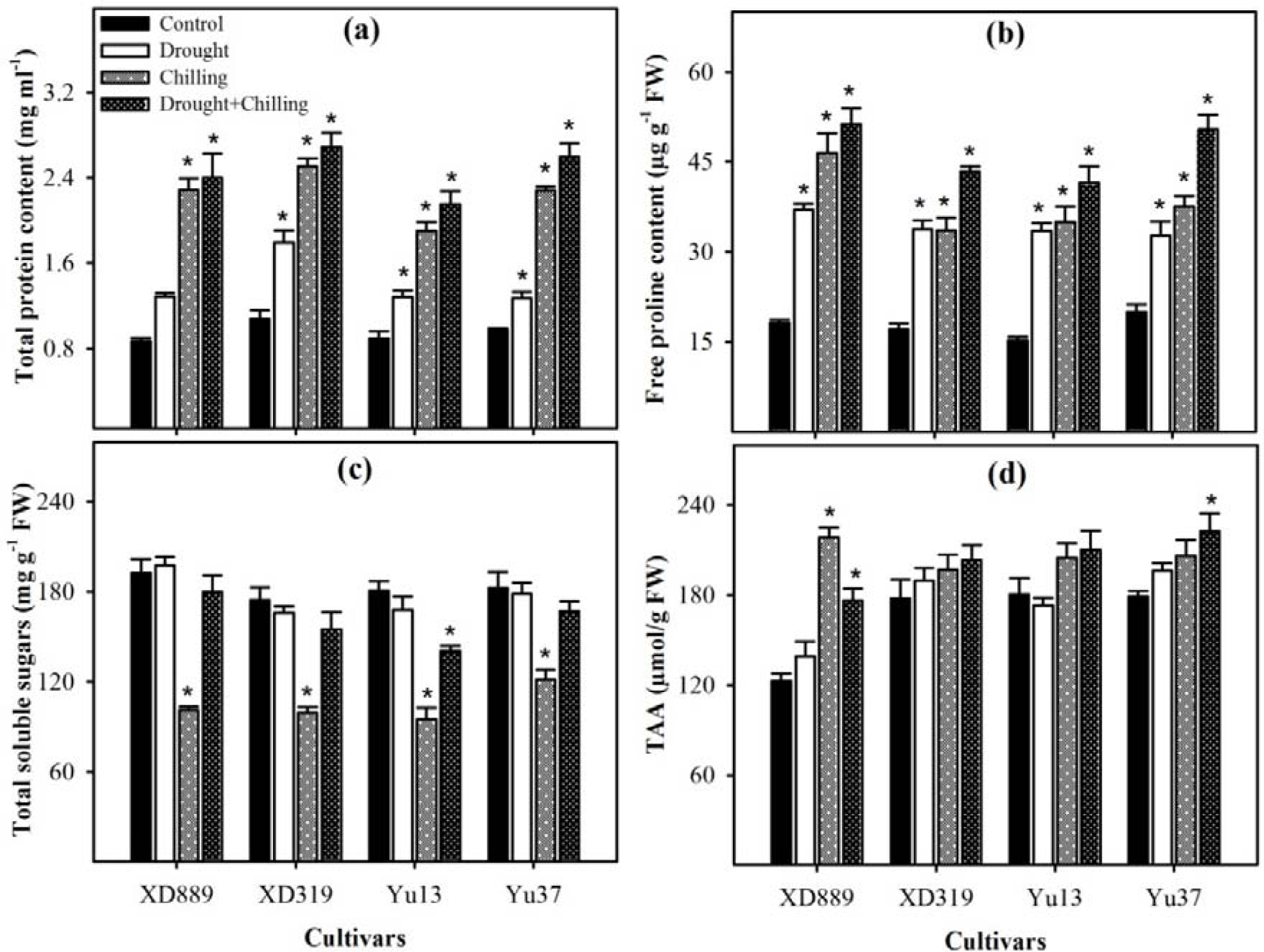
Accumulation of total soluble proteins (a), free proline (b), total soluble sugars (c), and total amino acid content (d) in the leaves of four maize cultivars as influenced by the drought, chilling and drought + chilling stress. Vertical bars above mean indicate standard error of three replicates. Mean value for each treatment with * indicate significant differences by the LSD-test (P ≤ 0.05).

## Discussion

Water deficit and suboptimal temperature are adverse environmental factors that impose drastic effects on crop plants [2,22]. Maize is comparatively more sensitive to water scarcity and low temperature conditions at early growth stages than later [4,31,32,33].

Present study indicated that maize shoot length, shoot fresh weight and shoot diameter was severely hampered by drought, chilling and combination of drought + chilling stresses (Figure 1 and 2). Compared with control, number of leaves and leaf width of maize were reduced under stress conditions (Figure 3), however, the negative effects of combined drought +chilling stresses were more severe for all these traits than their individual effects. Furthermore, the growth performance of hybrid maize was better than inbred cultivars under control as well as stress conditions (Figure 1,2 and 3). Early season drought coupled with chilling stress limited seed germination and early stand establishment that may be due to reduced intake of water during seed imbibition [4,18,19,21]. Drought and chilling-induced reductions in morphological traits of maize have also been reported previously [4,9,21]. Chilling stress may possibly reduce seedling growth by suppressing cell division, elongation in apical region and/or by causing physiological injuries in plant tissues [9,34], whereas, drought stress causes substantial reductions root and shoot growth and disturbs plant-water-relations [4,35].

Drought, chilling and drought + chilling stress reduced root length, root fresh weight, root surface area, root volume, total root length as well as root average diameter (Figure 4, 5 and 6), however, the root length of XD889 as well as root average diameter of XD889 and Yu37 were increased under water deficit conditions (Figure 4a, 6d). Mild drought conditions could enhance root length for water uptake whilst severe drought could severely reduce it. Previously, increased root length under drought stress was also recorded in sunflower and *Catharanthus roseus* [8,35].

Our results revealed that Chl a, Chl b and total Chl contents were severely affected in maize by individual and concurrent drought and chilling stresses (Figure 7), nonetheless, the Chl a and total Chl contents were increased in XD889 under chilling stress (Figure 7a and 7c). Previously, drought stress substantially reduced the leaf RWC and chlorophyll contents of the Xida 889 and Xida 319 cultivars which hamper the photosynthetic efficiency and plant growth of the Xida 889 and Xida 319 [36] Partelli et al. (2009) reported 30% and 27% reduction in Chl a and Chl b contents in coffee plants when day/night temperature was lowered from 25/20°C to 13/8°C, respectively [37]. Drought and chilling stress could lead to excessive ROS production which deteriorate chlorophyll contents in plant [38,39].

Drought, chilling and drought + chilling stress enhanced O^2□^, H_2_O_2_, OH^□^ and MDA contents in all maize cultivars Moreover, the level of ROS and MDA was higher under drought + chilling stress than their individual effects (Figure 8). In general, when the rate of ROS production exceeds the anti-oxidant enzyme activities then caused damage to essential cellular components [3,40]. Guo et al. (2006) reported that drought and chilling temperature increased the H_2_O_2_ contents in different rice cultivars [20]. The MDA contents is an indicator of extent of lipid peroxidation under stress conditions [17,18, 41]. The exposure of plants to unfavorable environmental conditions cause lipid peroxidation due to over-production of ROS [11,18].

Plants have developed a complex antioxidative defense system, consist of non-enzymatic and enzymatic components to overcome the excessive production of ROS [18,19, 42], however, the balance between ROS generation and antioxidant enzyme activities is critical in all plant species under stress conditions [12, 43]. Among various anti-oxidants, the SOD involves in the dismutation of O^•− 2^ to O_2_ and then H_2_O_2_, whilst CAT, POD, and GPX catalyze H_2_O_2_ to H_2_O and O_2_ [11,19,36]. In present study, antioxidant enzymes i.e., SOD, POD, CAT, GSH-PX, GR and GST activities were triggered under the drought, chilling and chilling + drought stress conditions (Figure9, 10). The contents of SOD and POD decreased under drought, chilling and drought + chilling stress conditions (Figure 9a and c) which was consistent with over-production of ROS and poor growth of maize seedlings in these treatments. Recently, some reports proved that the extreme stress conditions could imbalance the active oxygen metabolism as well as SOD and POD activities [17, 44, 45]. Furthermore, Cao et al. (2011) found that the SOD and POD activities were initially increased under drought and low temperature conditions and were decreased later with their prolonged exposure in oil palm [44]. Similarly, the concentration of CAT and GR were increased in different wheat cultivars under drought and low temperature stress [45,47,48]. According to present results, the activities of CAT and GR were also influenced by drought, chilling and drought + chilling stresses but response of maize cultivars were variable with stress conditions (Figure 9b and 9d). Sharma and Dubey (2005) reported substantial reduction in CAT activity in rice seedlings under drought stress [49]. Most often, stress-caused reduction in in the rate of protein turnover also responsible for reduction in CAT activity [42].

The level of GSH-PX and GST was up-regulated in all maize traits under drought, chilling and drought + chilling stress conditions (Figure 10a and 10b). Increased GR activity enhances GSH contents and is important to induce stress tolerance in plants [50]. Some GSTs function as GPXs to cleanse the products of oxidative stress. However, GPX improved tolerance against oxidative stress caused by salinity, drought and chilling stresses in transgenic *Arabidopsis*, whereas improved growth in tobacco seedlings was related to the overexpression of GST and GPX under various environmental stresses [51].

In addition, non-enzymatic antioxidants like GSH, vitamin C (Vc) and vitamin E (V_E_) were also been found crucial for stress tolerance in crop plants [11,19]. For instance, the GSH can directly detoxify the O^2•−,^ •OH, H_2_O_2_ and therefore, contributes in stress tolerance in plants [18,42,43, 52]. Our results showed that the V_E_ was reduced in Yu 13 under chilling and drought + chilling stress, whereas the GSH contents were reduced in XD889 and XD319 under drought and in Yu 13 under all stress conditions (Figure 9). Previously, Guo et al. (2006) stated that the GSH contents were increased under chilling stress but decreased under drought stress in different rice cultivars [20]. Hussain et al. (2016b) reported that the Vc and V_E_ are involved in various metabolic processes as non-enzymatic scavenger to quench ROS under stress conditions [19]. Moreover, the total antioxidant capability (T-AOC) of maize seedlings was up-regulated by drought, chilling and drought + chilling stress conditions (Figure 11d). The imbalance between ROS i.e., O^2□,^ H_2_O_2_ and OH^□^ in relation to antioxidants (Figures 9-11) might be resulted in oxidative stress which severely hampered the seedling growth under sub optimal temperature and low moisture supply. In general, the content of enzymatic and non-enzymatic scavengers triggered under stress conditions, however their levels were not enough to overcome the excessive ROS production. Drought, chilling and drought + chilling stresses triggered the production and accumulation of metabolites in all maize cultivars. Furthermore, the concentration of protein, proline and total amino acid (TAA) considerably increased in maize seedlings under drought, chilling and drought + chilling combine stress but the total amino acids (TAA) were reduced under drought stress in Yu13 (Figure 12). On the other hand, total soluble sugars were reduced in maize cultivars under stress conditions, but only increased in XD889 under drought stress, however, the total soluble sugars were lower under chilling stress than the drought and drought + chilling in all maize cultivars (Figure 12c). However, Krasensky and Jonak (2012) stated that the accumulation of numerous metabolites i.e., proline, carbohydrates, protein and amino acids are generally involved in maintaining cell turgor by osmotic adjustment, and sustaining redox metabolism to quench ROS [14]. Higher concentrations of proline were also found in oil palm under drought and low temperature condition [44]. Furthermore, compatible solutes protect the plants from osmotic stress by not posing any detrimental effects on enzymes, membranes, and other macromolecules even at higher concentrations [53,54].

